# Targeting the Mannitol Biosynthesis Pathway in *Aspergillus fumigatus*: Characterisation and Inhibition of Mannitol-2-Dehydrogenase

**DOI:** 10.64898/2026.06.15.732321

**Authors:** Stephanie Nguyen, Isis Pinner, C. Ruth Wang, Tara L. Pukala, Blagojce Jovcevski, John B. Bruning

**Author notes:** **Corresponding Authors:** Institute of Photonics and Advanced Sensing (IPAS), School of Biological Sciences, Adelaide University, South Australia, 5005, Australia, (B.J), (J.B.B). Australian Proteome Analysis Facility (APAF), Macquarie University, NSW 2109, Australia.

## Abstract

Infections caused by the opportunistic fungal pathogen *Aspergillus fumigatus* pose a serious public health system burden. The inherent limitations in existing antifungal drugs in conjunction with a rising emergence of antifungal resistance emphasizes an urgent need to identify and target alternative pathways crucial to survival and virulence. Targeting the fungal mannitol biosynthesis enzymes provides a promising avenue in the development of new antifungals due to the multifaceted roles mannitol fulfils in the fungal life cycle. However, a distinct lack of available structural information for these enzymes has hindered drug discovery efforts. We report the first crystal structure of mannitol-2-dehydrogenase from *A. fumigatus* in an unbound monomeric state (1.8 Å) and bound to its co-factor NADH (2.1 Å), via. a large, central cavity lined with positively charged residues that readily accommodates NADH. This interaction is further stabilised by a network of hydrogen bond interactions and π-π stacking between Phe45 and the nicotinamide ring of NADH. Furthermore, rigorous kinetic characterisation of *A. fumigatus* mannitol-2-dehydrogenase demonstrates the dose-dependent inhibitory activity of 1,4-benzoquinone, a cysteine-modifying small molecule inhibitor (IC_50_ = 1.2 ± 0.2 nM). In addition, intact MS and proteomic analysis further reveal that 1,4-benzoquinone modifies up to five cysteine residues of mannitol-2-dehydrogenase and displays antifungal activity against *A. fumigatus*, which is enhanced in combination with a front-line antifungal voriconazole. From this work, we have established the foundations for a novel antifungal drug discovery avenue that targets the fungal mannitol biosynthesis pathway to better treat aspergillosis and related pathogenic infections.

## Introduction

Aspergillosis is an infection commonly contracted by immunocompromised patients and is caused primarily by the pathogenic fungus *Aspergillus fumigatus*. Spores produced by *A. fumigatus* are ubiquitously found in both indoor and outdoor environments and are constantly inhaled into the respiratory system (1). However, they are neutralised in healthy human hosts by alveolar macrophages and neutrophils to avoid infection (2). Patients who have an impaired immune system, caused by chemotherapy, corticoid therapy, viral infection or organ transplant patients who require immune-suppressing drugs, become highly susceptible to the opportunistic nature of fungal pathogens ^(1)^. Infections caused by *A. fumigatus* are localised to the lung tissue, as observed in patients with allergic bronchopulmonary aspergillosis (ABPA), but can progress to more severe forms such as invasive aspergillosis which can affect multiple organs (1). Despite the availability of existing antifungal drugs used to treat aspergillosis, the mortality rate associated with these infections remains unacceptably high (3). This is attributed to a myriad of inherent limitations of these drugs including high toxicity, poor bioavailability in target tissues, poor activity spectrum and unfavourable drug interactions (3). These issues are further compounded by the growing rate of antifungal drug resistance. As the number of immunocompromised patients is expected to rise and the efficacy of existing antifungal drug treatment continues to decrease, there is a high demand for new classes of antifungals that target novel pathways to effectively treat human fungal infections. Targeting enzymes within the mannitol biosynthesis pathway has been proposed as a promising approach.

Mannitol is a six-carbon polyol that has been identified in various fungal structures including mycelia, fruiting bodies and conidia (4). Biosynthesised primarily through the activity of mannitol-2-dehydrogenase (M2DH, UniProtKB Q4WQY4) and mannitol-1-phosphate 5-dehydrogenase (M1P5DH, UniProtKB Q4X1A4), mannitol acts as a carbohydrate reserve, store of reducing power, osmolyte, and quencher of reactive oxygen species (ROS), all of which are roles that contribute to fungal survival and pathogenicity. Mannitol has been previously implicated in fungal survivability, specifically in the high stress tolerance of conidia (5–7). Disruption of mannitol production via knockout of the gene encoding M1P5DH in *Aspergillus niger*, a closely related species to *A. fumigatus*, conidia results in increased sensitivity to high temperatures, oxidative stress, freezing and lyophilisation (6). Thus, mannitol has been suggested to provide a protective effect against stressful conditions.

Studies into the role of mannitol in establishing virulence have also been explored in two human fungal pathogens, *Cryptococcus neoformans* and *A. fumigatus*. Both fungal species have been shown to produce and secrete mannitol during infection in an experimental disease model (8, 9). In rabbits with experimental meningitis inoculated with *C. neoformans*, higher levels of mannitol have been detected in cerebrospinal fluid (9). Similarly, rats with experimental aspergillosis showed an increase in mannitol levels in liver tissue and serum (8). To determine the effect of secreted fungal mannitol in these disease models, a low mannitol producing isolate of *C. neoformans* was injected intravenously into mice. In this experimental disease model, 100% of mice infected with low mannitol producing isolates survived 60 days post challenge whereas 100% of mice infected with wild-type *C. neoformans* H99 succumbed to the infection by 51 days. The severely diminished production of mannitol rendered the isolate 5,000 times less virulent than wild-type (5). Phenotypic characterisation of this mutant showed that it was also less resistant to environmental stresses such as temperature and salinity and was hypersensitive to ROS (10).

These studies suggest that mannitol plays a major role in fungal survival and pathogenicity in animal hosts. In addition, there are no human homologues of these mannitol biosynthesis enzymes and thus they have been suggested to be attractive antifungal drug targets (11). In this study, we present the first crystal structure of M2DH from *A. fumigatus*, in an unbound state and bound to NADH. These structural data have been used to provide insights into the active site residue composition, describe the characteristics of targetable substrate binding pockets and identify key interactions that stabilise the binding of NADH to the M2DH active site. The kinetic parameters of the interconversion reaction catalysed by M2DH are also described and we have demonstrated the inhibitory activity of 1,4-benzoquinone (1,4-BQ) against *A. fumigatus* M2DH. Together, these data provide the necessary foundations to establish an antifungal drug discovery strategy that specifically targets the mannitol biosynthesis pathway of pathogenic fungi.

## Materials and Methods

### Expression and Purification of Mannitol-2-Dehydrogenase

M2DH was expressed using an M2DH-pMCSG9 expression vector (Supplementary Methods) in *E. coli* BL21 (λDE3) cells were cultured in Luria broth supplemented with ampicillin (0.2 mg/mL) at 37 °C to an OD_600_ of 0.8. Protein expression was then induced for 16 h at 16 °C with the addition of 0.5 mM IPTG. Cells were harvested by centrifugation, resuspended in Buffer A (20 mM Tris-HCl pH 8.0, 500 mM NaCl, 10 mM imidazole, 2 mM β-mercaptoethanol), and stored at -80 °C until purification. Cells were thawed, then lysed in a M110 L microfluidizer processor (Microfluidics) and clarified by centrifugation before loading onto a 5 mL Zetasep Nickel NTA column equilibrated in Buffer A. The column was washed with 6 CV of a 90:10 ratio of Buffer A: Buffer B (20 mM Tris pH 8.0, 500 mM NaCl, 250 mM imidazole, 2 mM β-mercaptoethanol) to remove contaminants and M2DH was eluted using an imidazole gradient from 10 mM to 250 mM. The purity of M2DH was analysed using SDS-PAGE. Fractions containing M2DH were pooled, dialysed at 4 °C for 16 h into storage buffer (50 mM Tris-HCl pH 8.0, 0.5 mM EDTA, 5% glycerol, 1 mM DTT) and concentrated to 28.6 mg/mL in a 10,000 MWCO Amicon Ultra-15 Centrifugal Filter (Millipore). Aliquots of M2DH were flash frozen in liquid nitrogen and stored at -80 °C.

### Crystallisation of Mannitol-2-Dehydrogenase and Data Collection

M2DH microcrystals that formed in 30% PEG 4000, 0.2 M MgCl_2_, 0.1 M Tris pH 8.5 using sitting drop vapour diffusion were crushed and diluted 1:100 in reservoir solution to produce a seed stock. Crystals of unbound M2DH were grown using a modified random microseed matrix screening method (12). Using hanging drop vapour diffusion, unbound M2DH crystals formed in a 0.1:1:1 ratio of seed stock, M2DH (28.6 mg/mL) and 25% PEG 3350, 0.15 M Tris pH 8.5, 0.2 M NaCl. Cubic crystals (100 µm x 100 µm) of unbound M2DH were fully formed after two weeks of incubation at 16 °C. NADH-bound M2DH crystals were obtained by adding 1 µL of NADH solution (30 mM) dissolved in 27% PEG 4000, 0.22 M lithium sulphate, 0.1 M Tris pH 8.5 directly to unbound M2DH crystals, formed in 25% PEG 4000, 0.25 M lithium sulphate, 0.1 M Tris pH 8.5 and soaked for 17 h. Using a nylon CryoLoop, a single crystal was mounted, cryoprotected in *Paratone-N* (Hampton Research) and flash frozen in liquid nitrogen. Diffraction data of unbound M2DH crystals were collected at the Australian Synchrotron Macromolecular Crystallography Beam Line MX1 using an oscillation angle of 1° (yielding 360 frames) at a wavelength of 0.954 Å (^13^). Diffraction data of NADH-bound M2DH crystals were collected at the Australian Synchrotron Macromolecular Crystallography Beam Line MX2 using the Australia Cancer Research Foundation (ACRF) detector (14). Data was collected using an oscillation angle of 0.1° (yielding 3600 frames) at a wavelength of 0.954 Å.

### Data Processing and Structure Refinement

Data integration was performed using *XDS*, converted to an mtz file using *Pointless*, then scaling and merging was performed using *Aimless* (CCP4 program suite) (15–17). Molecular replacement was used to solve the phase problem using *Phaser MR* (CCP4 program suite) and M2DH from *Pseudomonas fluorescens* as an initial search model (1LJ8) (18). The solved structure of *A. fumigatus* M2DH (7RK4) was then used as a search model for the NADH-bound *A. fumigatus* M2DH structure. Manual rebuilding was completed in *WinCoot* and structure refinement in *Phenix.refine* (19, 20). Structure validation was assessed using *MolProbity* (21). Statistics of data processing and structure refinement are summarised in Table S1. Protein structure visualisation and figure preparation was completed using *PyMOL* version 2.3.4. Assessment of protein interfaces and domain architecture was completed using *PDBe PISA* version 1.52 and *InterPro*, respectively (22, 23). Coordinate files and structure factors of unbound M2DH and NADH-bound M2DH structures have been deposited in the Protein Data Bank under the following accession codes: 7RK4, 7RK5.

### Native Mass Spectrometry

The quaternary structure of M2DH was examined using a Synapt G1 HDMS (Waters) using a nanoelectrospray ionisation source (24). M2DH was buffer exchanged into 200 mM ammonium acetate (NH_4_OAc) (pH 6.8) using an Amicon Ultra-Centrifugal Filter (10,000 MWCO) at 4 °C. Protein concentration was adjusted to 15 µM and 2 µL of protein was loaded onto platinum-coated borosilicate glass capillaries prepared in-house. Key instrument parameters were as follows: capillary voltage (kV): 1.60; sampling cone (V): 50; extraction cone (V): 1.5; trap/transfer collision energy (V): 20/15; trap gas (L/hr): 5.5; backing gas (mbar): ∼4.5. Data analysed using MassLynx (v4.1, Waters) with minimal smoothing applied to spectra.

### Analytical Size-exclusion Chromatography

The average oligomeric size of M2DH (50 µM) was determined by analytical size-exclusion chromatography (SEC) using a Superdex 200 10/300 GL analytical-SEC (GE Healthcare) using methods described previously (24).

### Spectrophotometric Enzyme Assay for Kinetic Characterisation

Continuous activity assays of M2DH were performed as described previously with modifications (25). Assay mixtures contained 100 mM Tris-HCl pH 8.5 and saturating concentrations of substrates (500 mM mannitol and 4 mM NAD^+^ for the mannitol biosynthesis direction and 500 mM fructose and 1 mM NADH for the fructose biosynthesis direction). All assays were performed at 37 °C and the reaction was monitored using a PHERAstar FSX microplate reader (BMG LabTech) at 340 nm. The molar absorptivity of NADH (Δ_340_ = 6.22 mM^-1^ cm^-1^) was used to calculate the formation of NADH in the mannitol biosynthesis direction or the consumption of NADH in the fructose biosynthesis direction. To determine the Michaelis-Menten binding constants (K_M_), the concentration of mannitol (7.80 – 500 mM), NAD^+^ (0.0625 – 4 mM), fructose (7.80 – 500 mM) or NADH (0.125 – 1 mM) was varied whilst maintaining remaining substrates at saturating conditions. The reaction was started by adding M2DH to a final concentration of 12.5 nM, except in the determination of the NADH K_M_ which required a lower final concentration of 6.22 nM. The kinetic parameters were determined using data from three biological replicates plated in technical triplicate. Data was fitted to the Michaelis-Menten equation (GraphPad Prism 9).

### Spectrophotometric Enzyme Assay to Assess Inhibitory Activity of 1,4-Benzoquinone

Methods used to assay the inhibitory activity of 1,4-BQ against M2DH were based on methods used for kinetic characterisation, described above. Assay mixtures contained 100 mM Tris-HCl pH 8.5, 6.22 nM M2DH and 0.73 mM NAD^+^ (equal to the experimentally determined K_M_ of NAD^+^). Inhibitory assays were performed at 37 °C and the reaction was monitored using a PHERAstar FSX microplate reader (BMG LabTech) at 340 nm. The molar absorptivity of NADH (Δ_340_ = 6.22 mM^-1^ cm^-1^) was used to calculate the formation of NADH in the mannitol biosynthesis direction. To determine the half maximal inhibitory concentration (IC_50_), the concentration of 1,4-BQ (0.01 – 3.13 µM) was varied. The reaction was started by adding mannitol to a final concentration of 500 mM. The kinetic parameters were determined using data from three replicates plated in biological triplicate. Data was fitted to a dose-response curve (GraphPad Prism 9).

### Intact Mass Spectrometry Analysis

Apo-M2DH was diluted in 30 % (*v/v*) aqueous acetonitrile and 1 % (*v/v*) formic acid to a final protein concentration of 20 µM. Half of the apo-M2DH sample was reserved, and the remaining sample was then incubated with a 20-fold molar excess of 1,4-BQ in 4 % DMSO at 25 °C for 3 h. Apo- and 1,4-BQ-treated M2DH samples were stored at -20 °C until further analysis. Denatured protein mass spectra of both apo- and 1,4-BQ-treated M2DH were obtained using a 1260 LC system coupled to a 6230 TOF mass spectrometer (Agilent Technologies). Sample (5 µL) was introduced into the instrument by ESI-MS at a flowrate of 0.4 mL/min using 50 % (*v/v*) aqueous acetonitrile and 0.1 % (*v/v*) formic acid without chromatographic separation. The instrument conditions were set as follows: polarity set in positive mode; a mass range of 500 – 3200 *m/z*; capillary voltage (kV): 3.5; nozzle voltage (kV): 2; gas temperature (°C): 325. Mass spectra acquisition was performed using MassHunter software (vB.07.00, Agilent Technologies). Data analysis and deconvolution were performed using UniDec software (v.4.4.1) (26) with processing parameters as follows: charge range of 1 - 90; mass range of 50,000 to 65,000 Da; sampling of mass at every 10 Da; peak detection range set to 500 Da; peak detection threshold of 0.1.

### Antifungal Susceptibility Tests

*A. fumigatus Fresenius* spores (ATCC MYA-3626) were spread on Malt Extract Agar (MEA, Blakeslee’s formula) at 28 °C for 5 days. Spores were then resuspended in dH2O with 0.01 % *v/v* Tween20 to a concentration of 1.0 McFarland standard. The McFarland standard spore solution was applied to separate mannitol stripped MEA plates (100 μL total volume) and spread using six sterilised glass beads (Merck) per plate. Four sterile discs (6 mm, Whatman) were added to each plate (one in each quadrant), and plates were left to dry for 10 min. For each disc, 5 μL of test solution was added at a time. Each plate consisted of quadrants comprising: 1) dH_2_O (negative control); 2) voriconazole (500 ng total, positive control); 3) pBQ per plate (5–50 µg total); 4) pBQ (2.5–50 µg) combined with voriconazole (500 ng total). Samples containing pBQ were added last to minimise light exposure and plates were allowed to dry for a further 5 min. Sealed plates were covered to avoid light exposure and incubated at 37°C. *A. fumigatus* growth and inhibition was monitored at 48 h.

## Results

### Analysis of M2DH structure and oligomeric state

There has been a distinct lack of structural information for mannitol biosynthesis enzymes from fungal origins which has delayed the identification of targetable sites for antifungal drug discovery. To address this gap in knowledge, the X-ray crystal structure of M2DH from *A. fumigatus* in an unbound state (1.8 Å) was solved (Figure 1A-B). The structure is comprised of 22 α-helices, 16 β-strands, 10 3_10_-helices and 2 π-helices, arranged into two domains (Figure 1B). The N-terminal domain (residues 1-201) features a Rossmann-type fold, characterised by the alternating βαβ motif, which is commonly observed in cofactor-binding proteins whereas the C-terminal domain (residues 232-475) is predominantly composed of α-helices. The structure of M2DH solves as a crystallographic dimer in the asymmetric unit, forming a protein-protein interface encompassing a surface area of 879 Å^2^ (Figure 1A). According to the Complex Formation Significance Score calculated by *PDBe PISA*, this dimeric interface is likely to be the result of crystal packing. To validate these results, analytical-SEC and native MS was used to determine the oligomeric state of *A. fumigatus* M2DH (Figure 1C-D). These data show the presence of a dominant species that corresponds to one monomeric unit of M2DH (measured molecular weight: 57,465 Da, theoretical molecular weight: 57,442 Da) with a small proportion of dimeric species also observed (Figure 1C). This dimer formation under native MS conditions is mediated by transient hydrophobic and electrostatic interactions at dimer interfaces that are artificially stabilised in the gas phase. Subsequent analysis of oligomeric state by analytical-SEC of M2DH shows a single peak elution at 15.2 mL, indicative of monomeric M2DH in solution (Figure 1D, dashed line). Together, these data indicate that M2DH from *A. fumigatus* appears to be functional in a monomeric state.

**Figure 1:**
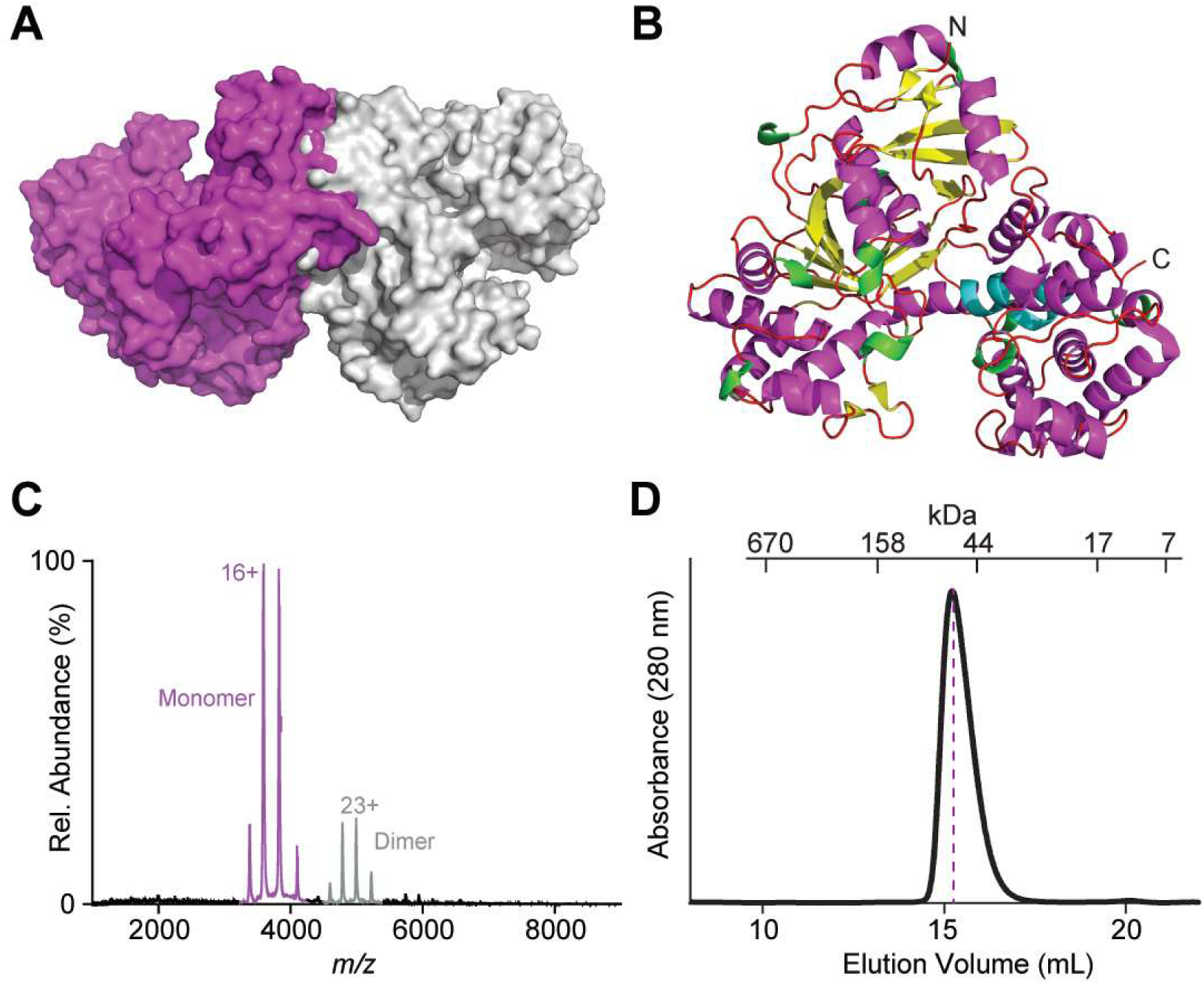
Crystal structure of a monomeric M2DH from *Aspergillus fumigatus.* **(A)** The structure of M2DH solves as a crystallographic asymmetric dimer due to crystal packing. **(B)** The structure of M2DH from in cartoon representation (1.8 Å, 7RK4). Secondary structure is colour-coded as follows: α-helix (magenta), β-strand (yellow), 3_10_-helix (green), π-helix (cyan) and loops (red). **(C)** Native MS shows M2DH (15 μM in 200 mM ammonium acetate, pH 6.8) is predominantly monomeric (purple) with a small population of dimer (grey). **(D)** Analytical-SEC of M2DH (50 μM) in 50 mM phosphate buffer (pH 7.4). Elution volumes of molecular mass standards are indicated above.

### High structural similarity exists between homologous enzymes

Consistent with its conserved role in mannitol biosynthesis, the structure of M2DH from *A. fumigatus* shows a high degree of sequence and structural similarity in comparison to M2DH from *P. fluorescens* in terms of its overall fold and domain architecture (Figure S2 & S3). Although these homologous enzymes are derived from different domains of life and exhibit moderate sequence conservation (44% sequence identity), superimposition of the α-carbon atoms that comprise the backbone residues reveals a highly conserved overall fold, represented by their low RMSD of 3.1 Å (Figure S3). This superimposition also indicates that the characteristic N- and C-terminal domains adopted by M2DH are also conserved between homologous enzymes (Figure S3).

### Characterisation of the NADH-binding site

The interconversion of mannitol and fructose, catalysed by M2DH, requires the use of an NAD^+^/NADH co-factor. Visualisation of the electrostatic surface of M2DH reveals a large, central cavity that is formed at the boundary of the N- and C-terminal domains. This substrate-binding site is predominantly lined by basic amino acid residues. The strong electron density observed in the central cavity of holo-M2DH supports modelling of the NADH co-factor (occupancy of 0.79 and 0.92 for each monomer in the asymmetric unit) (Figure 2A). The abundance of positive charges within the active site readily accommodates the negatively charged phosphate groups that bridge the adenosine and nicotinamide riboside groups of NADH (Figure 2A). The nicotinamide riboside moiety is positioned deeper into this binding pocket whereas the adenosine component of NADH resides within a more surface exposed ridge in a region that is largely neutral in electrostatic potential (Figure 2A). The stabilisation of NADH within this binding pocket is facilitated by π-π stacking interactions, an extensive hydrogen bond network and hydrophobic interactions. The pyridine ring of nicotinamide forms a π-π stacking interaction with the Phe45 sidechain whereas the amide forms hydrogen bonds (2.9 Å and 3.1 Å) with the Thr242 backbone (Figure 2B). Additional hydrogen bonds are established between the bridging phosphate groups and the backbone of Gly44 (2.9 Å) and Phe45 (2.8 Å), as well as between both hydroxyl groups of the ribose in the adenosine moiety and the side chain of Asp77 (2.4 Å and 2.9 Å) (Figure 2B).

**Figure 2:**
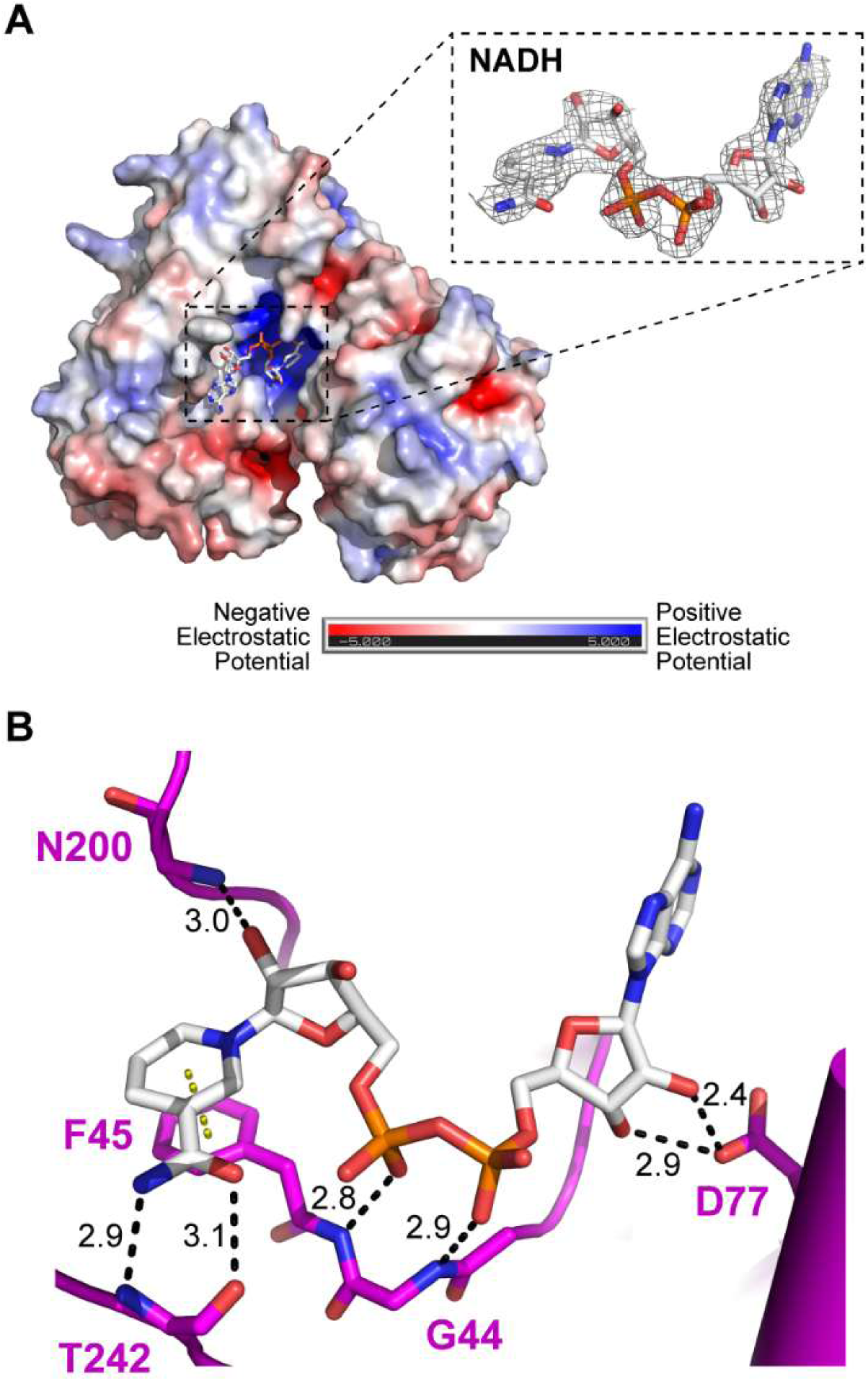
The reduced form of NADH bound to M2DH from *A. fumigatus*. **(A)** The electrostatic surface diagram shows NADH occupying a positively charged cavity. The simulated annealing composite omit map (2Fo-Fc) contoured at 1 σ supports the modelling of NADH (inset). **(B)** Extensive hydrogen bonding (black dashed lines) and π-π stacking (yellow dashed lines) between NADH (sticks) and binding site residues stabilise this interaction.

### Kinetic characterisation and inhibition of M2DH

M2DH catalyses the interconversion of mannitol and fructose, using either NAD^+^ or NADH as essential cofactors. The apparent Michaelis-Menten binding constants (K_M_) measured for both carbohydrate substrates, mannitol and fructose, are consistent with those measured in kinetic characterisation experiments conducted with *A. fumigatus* M2DH performed previously (Table 1) (Figure 3A-D) (11). However, the apparent K_M_ values measured for both cofactors, NAD^+^and NADH, recorded in this study were consistently higher by almost 7-fold and 11-fold, respectively. Calculations of the specificity constant ((*k_cat_/*K_M_) in s^-1^ mM^-1^) indicate that M2DH from *A. fumigatus* preferentially converts mannitol to fructose (*k_cat_*/K_M (mannitol)_ = 5.1 ± 0.55 s^-1^ mM^-1^) as opposed to the opposite reaction (*k_cat_*/K_M (fructose)_ = 0.71 ± 0.12 s^-1^ mM^-1^).

**Figure 3:**
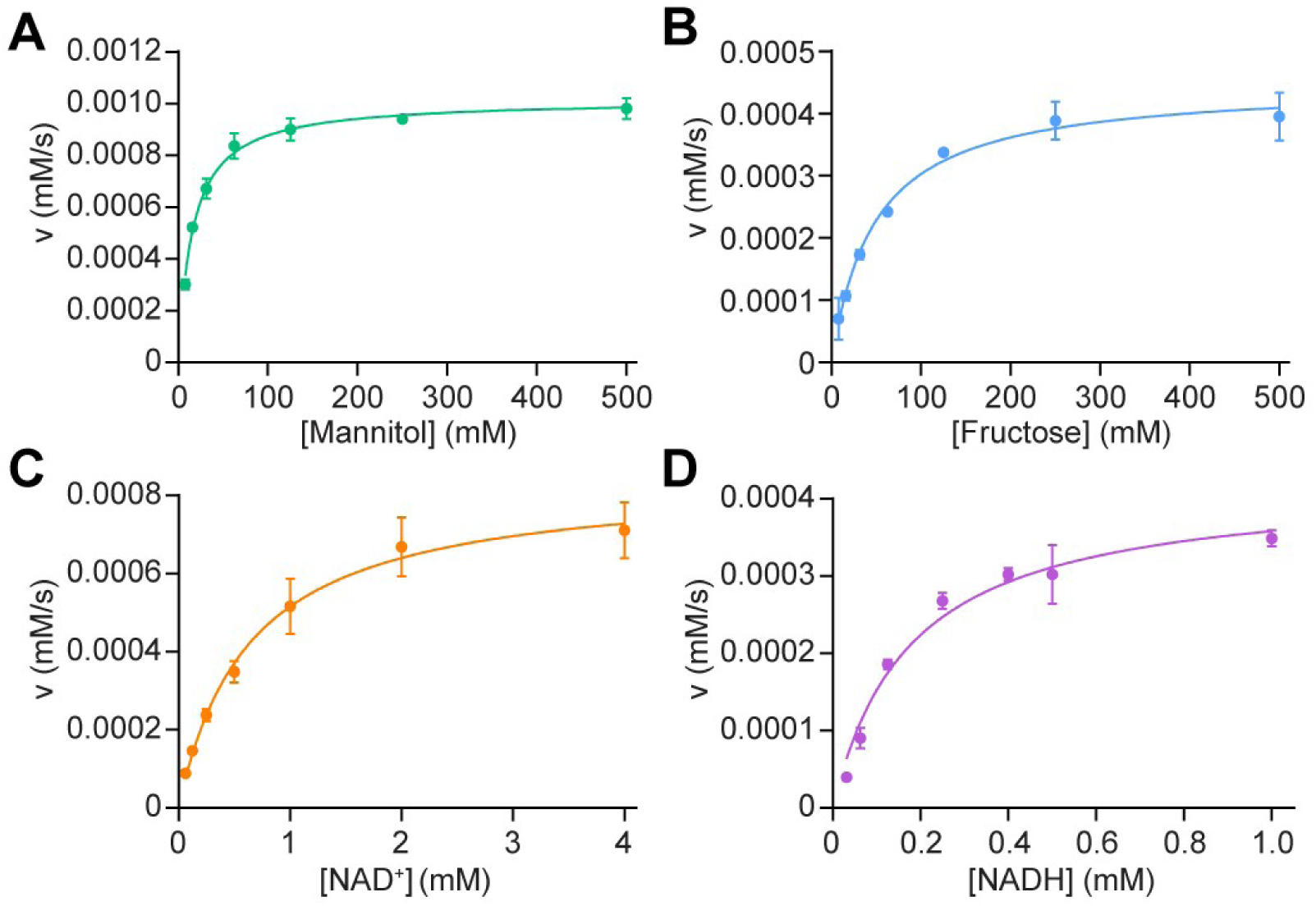
M2DH activity assays in the presence of various substrates. Representative steady-state M2DH activity assays fitted to the Michaelis-Menten equation to determine the Michaelis-Menten binding constant (*K_M_*) and catalytic constants (*k*_cat_) of the substrates, **(A)** mannitol, **(B)** fructose, **(C)** NAD^+^ and **(D)** NADH. Data presented as mean ± SD (n = 3).

**Table 1:**
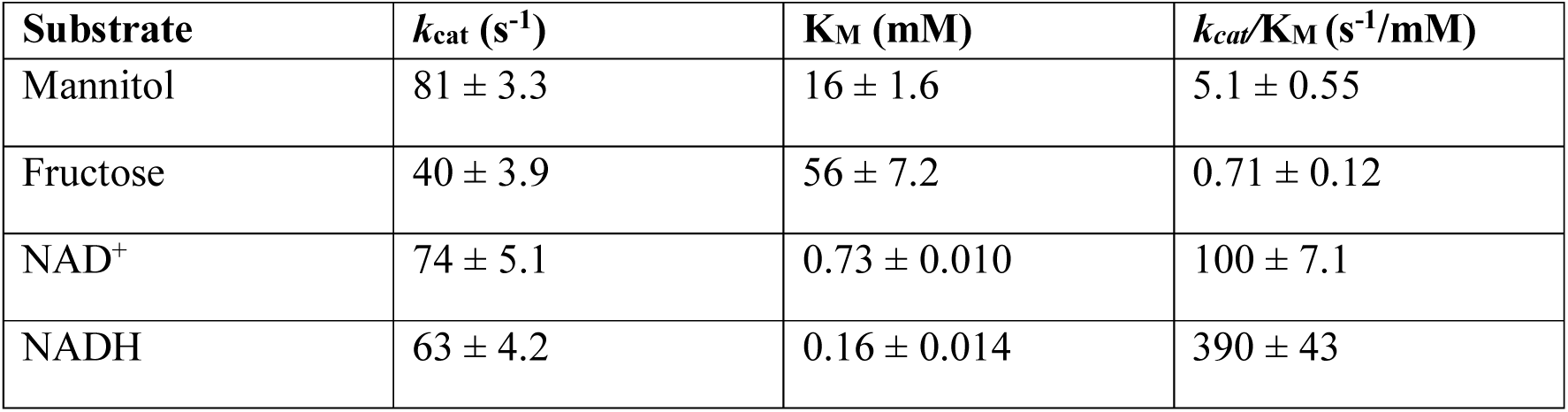
Apparent Michaelis-Menten constants (K_M_) and catalytic constant (*k*_cat_) of M2DH with substrates used in mannitol and fructose biosynthesis reactions. Data presented as mean ± SD (n = 3).

| Substrate | $k_{cat}$ ($s^{-1}$) | $K_M$ (mM) | $k_{cat}/K_M$ ($s^{-1}/mM$) |
| --- | --- | --- | --- |
| Mannitol | $81 \pm 3.3$ | $16 \pm 1.6$ | $5.1 \pm 0.55$ |
| Fructose | $40 \pm 3.9$ | $56 \pm 7.2$ | $0.71 \pm 0.12$ |
| NAD<sup>+</sup> | $74 \pm 5.1$ | $0.73 \pm 0.010$ | $100 \pm 7.1$ |
| NADH | $63 \pm 4.2$ | $0.16 \pm 0.014$ | $390 \pm 43$ |

### Mechanism of M2DH inhibition and antifungal activity of 1,4-benzoquinone

The small molecule compound, 1,4-BQ, was previously identified as a potent inhibitor of M2DH from *Acetobacter xylinum* (*K*_i_ = 0.18 mM) (27). Using an *in vitro* enzyme assay, 1,4-BQ was also shown to have dose-dependent inhibitory activity against *A. fumigatus* M2DH (Figure 4A). The half maximal inhibitory constant (IC_50_) determined was 1.2 ± 0.2 nM (Figure 4A). A potential mechanism in which 1,4-BQ inhibits M2DH activity is through thiol S-alkylation via Michael addition of cysteine residues, as observed previously for monoamine oxidases (28) and α-lactalbumin (29). A combined intact MS and bottom-up proteomics approach was utilised to determine whether 1,4-BQ modifies M2DH (Figure 4B-C, Figure S4). Intact MS analysis of apo M2DH showed populations of both full-length and cleaved methionine (-Met) M2DH (Figure 2B), which arises from proteolytic cleavage of the N-terminal methionine during recombinant protein expression by methionine aminopeptidase. Intact MS coupled with mass deconvolution analysis of 1,4-BQ treated M2DH identified mass shifts corresponding to four (+424 Da) and five (+503 Da) benzoquinone modifications on both full length and -Met M2DH (+108 Da) (Figure 3C). Bottom-up proteomic analysis of 1,4-BQ treated M2DH exhibited 76% sequence coverage and only covering one Cys residue (Cys69), due to decreased ionizability of Cys-modified peptides affecting their detection, where we observe this 1,4-BQ specific modification (+156 Da) on Cys69 (Figure S4). Together, with the bottom-up proteomics data, the data indicates that 1,4-BQ induces a series of thiol S-alkylation modifications on Cys residues, potentially inhibiting M2DH activity.

**Figure 4:**
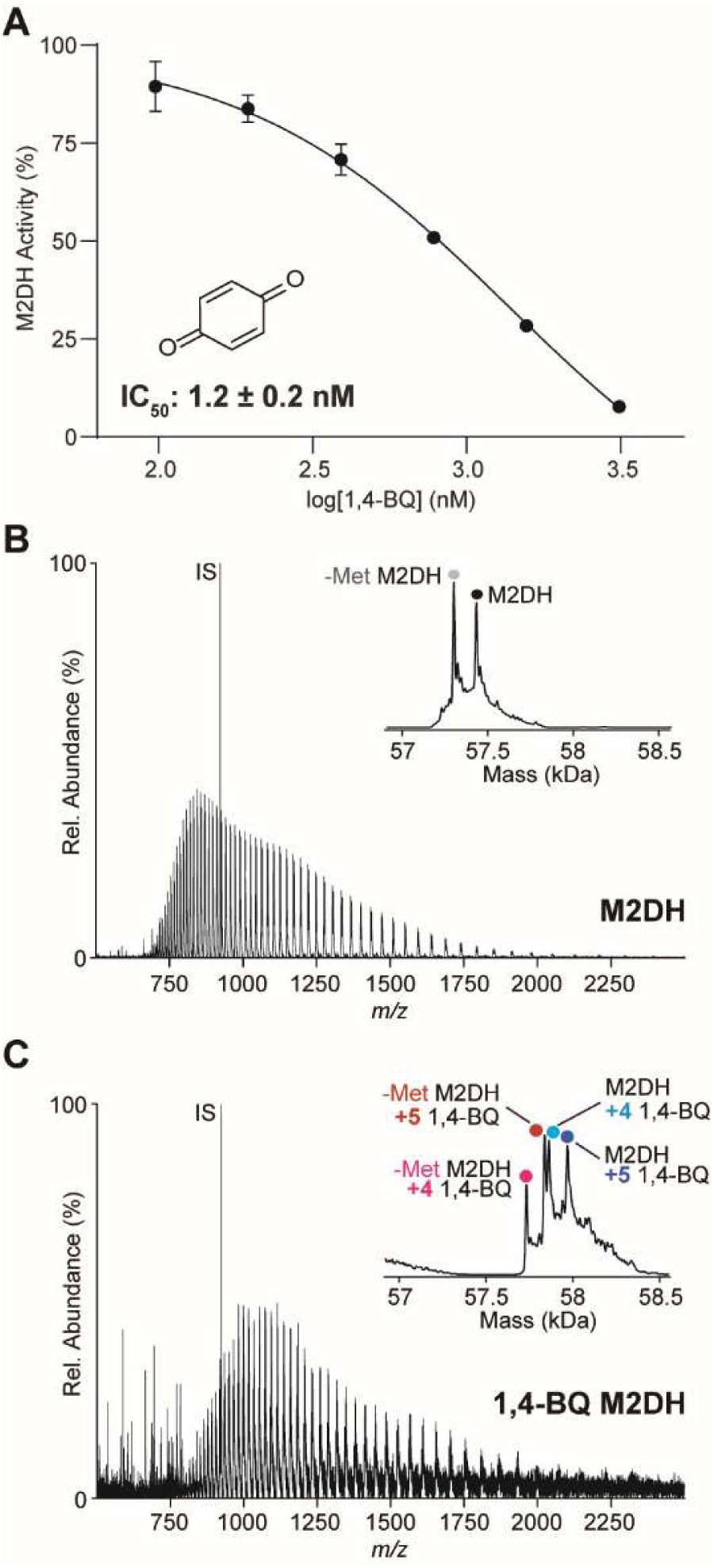
1,4-BQ inhibits *A. fumigatus* M2DH activity by residue modification. **(A)** 1,4-BQ induces dose-dependent inhibition of *A. fumigatus* M2DH activity *in vitro*. Data presented as mean ± SD (n = 3). **(B)** Intact MS and mass deconvolution (inset) of apo M2DH shows the presence of both cleaved methionine (-Met, grey) and full length (black) M2DH. **(C)** Intact MS and mass deconvolution (inset) exhibits mass shifts corresponding to four (pink and red) and five (cyan and blue) residues being modified by 1,4-BQ in both cleaved methionine and full-length M2DH (internal standard: IS in panel **B** and **C**).

The ability of 1,4-BQ to inhibit M2DH activity and thereby exert antifungal activity, in combination with known antifungals (e.g. voriconazole), was subsequently examined using disc diffusion assays. After 48 h incubation, the area of inhibition for the voriconazole was consistent at 500 ng measuring at a diameter of ∼15 mm (Figure 5), whereas the dH_2_O negative control showed no inhibition. The area of inhibition for 1,4-BQ showed dose response with inhibition being observed as low as 5 µg. Interestingly, combination of 1,4-BQ with voriconazole displayed increased inhibition of mycelia growth, closer to the centre of the disc (Figure 5). Together, this data shows that inhibition of M2DH activity by 1,4-BQ could potentially be rationalised by irreversible cysteine modification, thereby affecting *A. fumigatus* growth.

**Figure 5:**
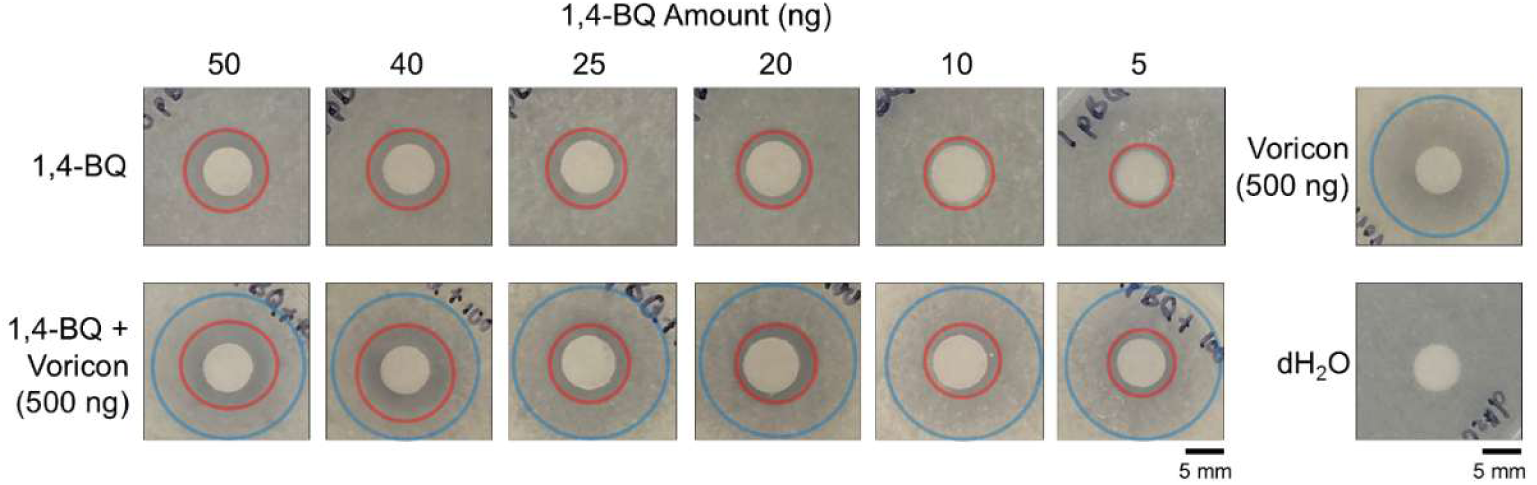
1,4-BQ inhibits *A. fumigatus* growth in combination with voriconazole. Disc diffusion assays monitoring *A. fumigatus* growth in the presence of a dilution series of 1,4-BQ alone (red) and in combination with voriconazole (Voricon, blue). All plates and conditions were repeated in at least two independent experiments. Scale bar: 5 mm.

## Discussion

Mannitol biosynthesis is an integral metabolic process undergone by several pathogenic fungi to survive normal environmental conditions and during the infection of a host (8, 9). When mannitol is produced in high abundance, it is able to efficiently fulfil its roles as a storage of carbohydrates, quencher of ROS, osmoregulator and regulator of pH (4). These multifaceted roles ultimately contribute to the survival and persistence of pathogenic fungi within a human host. A novel approach to developing drugs that are effective against a broad spectrum of pathogenic fungi lies in the inhibition of the mannitol biosynthesis enzymes. In the existing literature, there have been some interest in targeting two key enzymes capable of producing mannitol from this pathway, M2DH and M1P5DH (11). However, there has been a distinct absence of structural data which has impeded drug discovery efforts. In this study, we present the first crystal structure of M2DH from *A. fumigatus* to establish the necessary foundations for a drug discovery project poised to accelerate the development of novel inhibitors that target the fungal mannitol biosynthesis pathway.

The crystal structures from *A. fumigatus* M2DH unbound and bound to NADH were solved to a resolution of 1.80 Å and 2.10 Å, respectively (Figures 1 and S3). Analysis of the overall structure of *A. fumigatus* M2DH reveals a large degree of structural similarity, in terms of their overall fold and domain architecture, exists between homologous enzymes despite their low sequence similarity (Figure S2). Furthermore, the analysis of oligomeric interfaces in the crystal structures, analytical-SEC and native mass spectrometry data confirm that M2DH from *A. fumigatus* is monomeric. Due to the highly conserved nature of these enzymes, the dissemination of a publicly available structure of M2DH from *A. fumigatus* may be of interest to other research groups as it can serve as a preferable template for the modelling of homologous enzymes from pathogenic fungi. A subsequent application of these homology models would be in *in silico* screening experiments of large compound or fragment libraries to identify novel inhibitors. This approach would be effective for homologous enzymes that may not be amenable to crystallisation.

To identify small molecule inhibitors that are effective against M2DH, it is imperative to define the most ‘druggable’ binding site using this information to define the target site for high throughput *in silico* screening experiments. There is a singular, central cavity that lies between the N- and C-terminal domains of M2DH, capable of binding the substrates and co-factors required in the interconversion reaction (Figure 2A). Based on structural superimposition between the NADH-bound *A. fumigatus* M2DH structure and the NAD^+^- and D-mannitol-bound *P. fluorescens* M2DH structure, the co-factor and substrate bind closely to each other, each in a designated region of this pocket (Figure 6). Specifically, mannitol is positioned adjacent to the nicotinamide group of NAD^+^/NADH and extends deeper into the positively charged pocket. From the co-factor-bound structure of *A. fumigatus* M2DH, NAD^+^/NADH is stably held in the positively charged pocket by an extensive network of hydrogen bond and π-π-stacking interactions (Figure 2B). Closer inspection of this binding pocket shows a groove that extends above and below the adenine group of NADH into a solvent-exposed region (Figure 2A). Potential approaches to inhibitor design could explore analogues of NADH, particularly those that have chemical modifications on the adenine group to form additional interactions with surrounding residues in the solvent-exposed grooves. An alternate approach may involve joining an NADH and mannitol analogue using a short chemical, non-hydrolysable linker to exploit both regions of the binding pocket. This may be feasible due to the proximity of the two binding sites in the central cavity.

**Figure 6:**
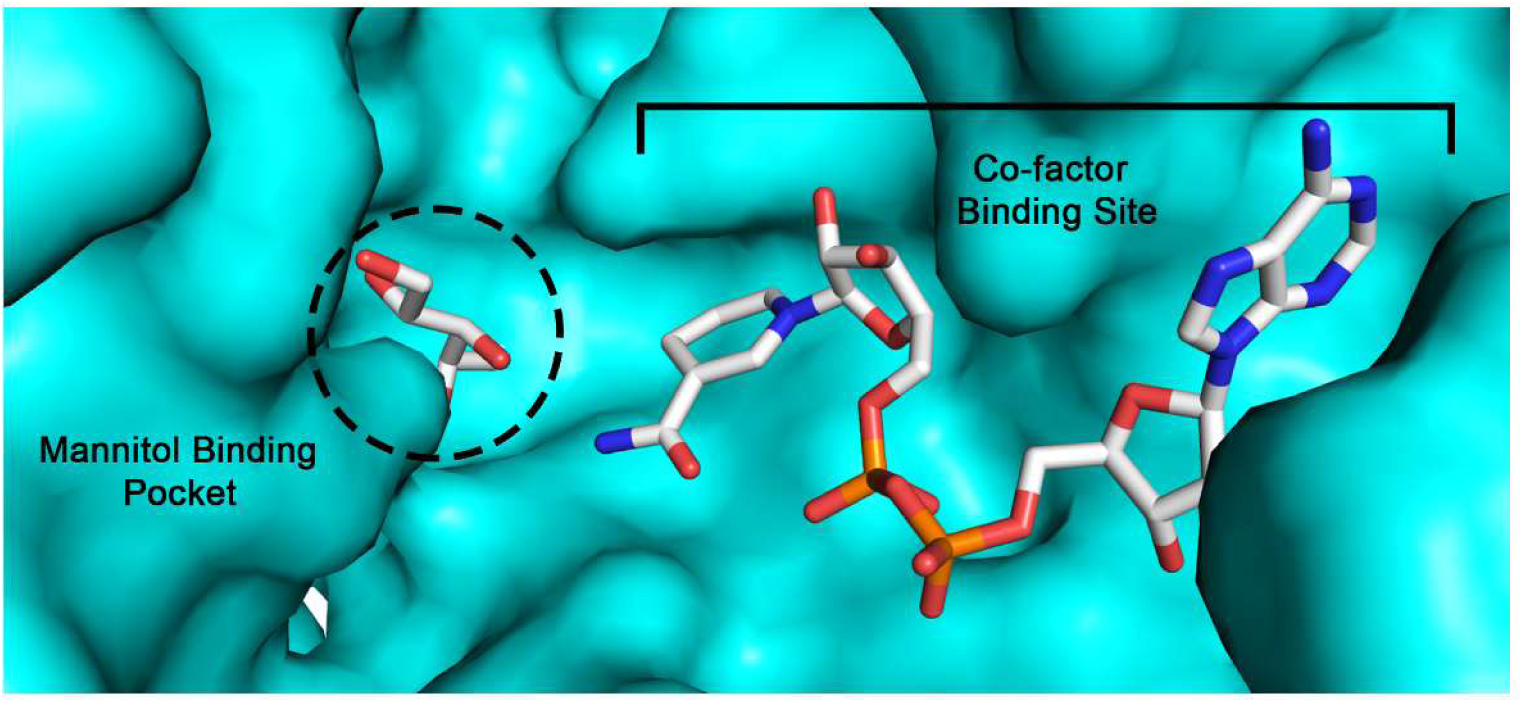
Structural superimposition of the NADH-bound *A. fumigatus.* M2DH structure and the NAD^+^- and D-mannitol-bound *P. fluorescens* M2DH structure reveals two distinct pockets located within the central cavity in which the substrates bind. The surface representation of *A. fumigatus* M2DH is coloured in cyan, the mannitol and NADH molecules are represented as sticks.

Kinetic characterisation studies were completed for all substrates used in the interconversion reaction (Table 1). Analysis of the specificity constant reveals a preference for M2DH to catalyse the turnover of mannitol to fructose, using NAD^+^ as a co-factor, as opposed to the opposite reaction. Although M2DH can produce mannitol, these kinetic data indicate that M1P5DH may instead be primarily responsible for mannitol biosynthesis whereas M2DH only plays a supporting role. This has implications for the targeted disruption of the mannitol biosynthesis pathway in pathogenic fungi by small molecule inhibitors. An effective approach may require simultaneous inhibition of M2DH and M1P5DH enzymes to downregulate mannitol production. Phenotypic characterisation of a *C. neoformans* mutant that produced low levels of mannitol revealed a significant reduction in virulence compared to wild-type and an increased sensitivity to environmental stressors (5, 10). Thus, dual targeting of the *A, fumigatus* mannitol biosynthesis enzymes may be sufficient to induce a comparable phenotype. As a result, this could increase fungal susceptibility to killing mechanisms employed by the immune system and resistance to changes in the host environment will be impaired.

As a proof of concept, we have shown that M2DH enzyme activity can be modulated using a small molecule inhibitor. We show that 1,4-BQ exhibited dose-dependent inhibition against M2DH from *A. fumigatus*. This compound has been shown to be effective against several enzymes, including monoamine oxidase, and its derivatives have been shown to have antimicrobial activity against *Staphylococcus aureus* and *Mycobacterium tuberculosis* (28, 30). Mechanism of action studies that have focused on compounds that feature a 1,4-BQ scaffold indicate that it modifies the free thiol group on cysteine sidechains, resulting in irreversible inhibition by functioning as a covalent inhibitor (28). Our MS data suggest that a similar mechanism elicits the inhibition of M2DH activity by 1,4-BQ through benzoquinone modifications of all M2DH cysteine residues. There are several structural analogues of 1,4-BQ in which chemical modifications have been added to the existing scaffold. Future work should explore the structure-activity relationships of these analogues as inhibitors of M2DH which inhibit *A. fumigatus* growth.

From this work, we have structurally and kinetically characterised the mannitol biosynthesis enzyme, M2DH from *A. fumigatus*. From these data, we have provided insights into targetable binding pockets to aid structure-guided drug design of novel inhibitors. As a proof of concept, we have shown that the small molecule 1,4-BQ can modulate the activity of M2DH. This work establishes a novel antifungal drug discovery avenue targeting the fungal mannitol biosynthesis pathway to combat aspergillosis and related infections.

## Accession Codes

M2DH: Q4WQY4

M1P5DH: Q4X1A4

## PDB Codes

M2DH: 7RK4

M2DH + NADH: 7RK5

## Acknowledgements

We thank Flinders Analytical (Flinders University, Australia) for access to native MS instrumentation. We also thank Lewis McFarlane (Walter Eliza Hall Institute, Melbourne) for his assistance with the purification of 1,4-benzoquinone. S.N., I.P. and C.R.W. are recipients of an Australian Government Research Training Program stipend scholarship. B.J. is supported by a Hospital Research Foundation Research Fellowship (2023/QA25313).

